# Oxidative stress reshapes diatom-microbiome interactions by shifting benefits from mutualistic to opportunistic bacteria

**DOI:** 10.64898/2026.06.18.733249

**Authors:** Nana Ankrah, Courtney Swink, Nathan McCall, Kristina Rolison, Christina Ramon, Peter K. Weber, Rhona K. Stuart, Xavier Mayali

## Abstract

The roles of reactive oxygen species (ROS) as signaling molecules and inhibitors of phytoplankton growth are well documented. While phytoplankton physiological mechanisms for ROS detoxification are well characterized, the role of heterotrophic bacterial partners in ROS alleviation and outcomes for these bacteria remain poorly understood. Here, we examined how extracellular hydrogen peroxide (H_2_O_2_) shapes nutrient exchange between the diatom *Phaeodactylum tricornutum* and two phycosphere bacteria. From an initial screen of 20 bacteria, we identified a “helper” (*Muricauda* sp.) that enabled *P. tricornutum* to survive acute H_2_O_2_ stress and a “non-helper” (*Algoriphagus* sp.) that did not. Using nanoscale secondary ion mass spectrometry (nanoSIMS), we tracked diatom-derived carbon and nitrogen (^13^C and ^15^N) transfer to each bacterial partner under ROS stress. Oxidative stress disrupted diatom metabolism and altered nutrient transfer: diatom-derived carbon and nitrogen incorporation was significantly reduced in the helper but increased in the non-helper under H_2_O_2_ stress. Growth assays revealed that the helper preferentially utilized exudates from healthy, intact hosts, whereas the non-helper did not grow on exudates but thrived on lysates from damaged or lysed cells. Together, these findings indicate the helper was better adapted to accessing resources from living hosts, while the non-helper relied on nutrients released through ROS-induced host damage. Our results highlight oxidative stress as a key driver of algal-bacterial interactions and suggest that bacterial resource-acquisition strategy underlies host protection: bacteria utilizing healthy-host exudates are more likely to protect hosts from oxidative stress, while those benefiting from host damage are not, despite retaining ROS detoxification capacity.

## Main text

The importance of reactive oxygen species (ROS) to phytoplankton is well documented. At low concentrations, ROS may act as signaling molecules and play roles in cell physiology, while at high concentrations they induce cellular damage via the degradation of biomolecules like proteins and nucleotides, ultimately leading to cell death (1). ROS are omnipresent in aerobic environments, formed by photochemical transformation of organic matter and as by-products of microbial metabolism (2, 3). To protect against ROS damage, many phytoplankton living in aerobic environments use enzymatic (e.g. catalase, superoxide dismutase) and non-enzymatic (e.g. carotenoids, reduced glutathione) mechanisms to scavenge ROS and maintain normal cellular homeostasis (4,5). These defenses notwithstanding, phytoplankton mortality increases under certain extreme conditions where the rate of ROS production exceeds the cellular scavenging capacity.

To enhance survival under extreme ROS conditions, some phytoplankton depend on and benefit from associations with ROS-degrading “helper” heterotrophic bacterial partners (6,7), but we have limited data on the reciprocal benefits heterotrophic bacteria gain from this interaction. Helper-beneficiary interactions have been most extensively documented in the marine cyanobacterium *Prochlorococcus*, which depends on co-occurring heterotrophs for protection against oxidative damage (7,8,9). Yet even for this system, our understanding of the benefits of ROS scavenging to the heterotrophic bacterial partner remains elusive.

We hypothesized that ROS stress alters phytoplankton-bacteria interactions by altering the form in which phytoplankton-derived nutrients are released, thereby differentially benefiting bacterial partners with symbiotic or opportunistic life strategies. Specifically, we predicted that (i) helper bacteria that mitigate oxidative stress would benefit primarily by maintaining access to phytoplankton exudates released by healthy intact cells, whereas (ii) non-helper bacteria would benefit disproportionately under oxidative stress by exploiting nutrients released through host cell damage and lysis. Under this framework, increasing oxidative stress should shift carbon and nitrogen transfer away from cooperative helper taxa and toward opportunistic non-helper taxa, even when both are capable of degrading reactive oxygen species in isolation.

To test these hypotheses, we used the diatom *Phaeodactylum tricornutum* and its phycosphere microbiota to investigate how external H_2_O_2_, a common ROS, impacts the transfer of phytoplankton-derived carbon and nitrogen to bacterial partners. We first separately incubated *P. tricornutum* with 22 members of its phycosphere microbiota to identify helper bacteria that were effective at providing protection against acute H_2_O_2_ stress and non-helper bacteria that did not (Figure S1). Bacteria that provided protection to *P. tricornutum* against H_2_O_2_ exposure relative to axenic cultures were classified as helper bacteria, and those that did were classified as non-helper. We identified two bacterial isolates, *Muricauda* (a helper bacterium) and *Algoriphagus* (a non-helper bacterium), for further experimentation.

While both *Muricauda* and *Algoriphagus* grew across a broad range of H_2_O_2_ concentrations in monoculture, *Muricauda* was more sensitive to high H_2_O_2_ concentrations than *Algoriphagus* (Figure S2). Despite this greater sensitivity, only *Muricauda* enabled *P. tricornutum* survival in coculture following H_2_O_2_ addition (Figure 1A). In coculture, *Muricauda* cultures exhibited greater reduction in extracellular H_2_O_2_ concentrations than *Algoriphagus* cultures under both moderate and high H_2_O_2_ conditions (Figure S3), suggesting that the ability to mitigate oxidative stress (rather than intrinsic H_2_O_2_ tolerance) may underlie *Muricauda*’s protective effect on *P. tricornutum*. These findings indicate that bacterial H_2_O_2_ tolerance and scavenging alone is insufficient to define helper status. This decoupling suggests that effective helper status likely depended on additional traits, such as spatial proximity to algal cells, the localization and regulation of ROS detoxification, or complementary metabolic interactions that stabilize host physiology under ROS stress.

**Figure 1.**
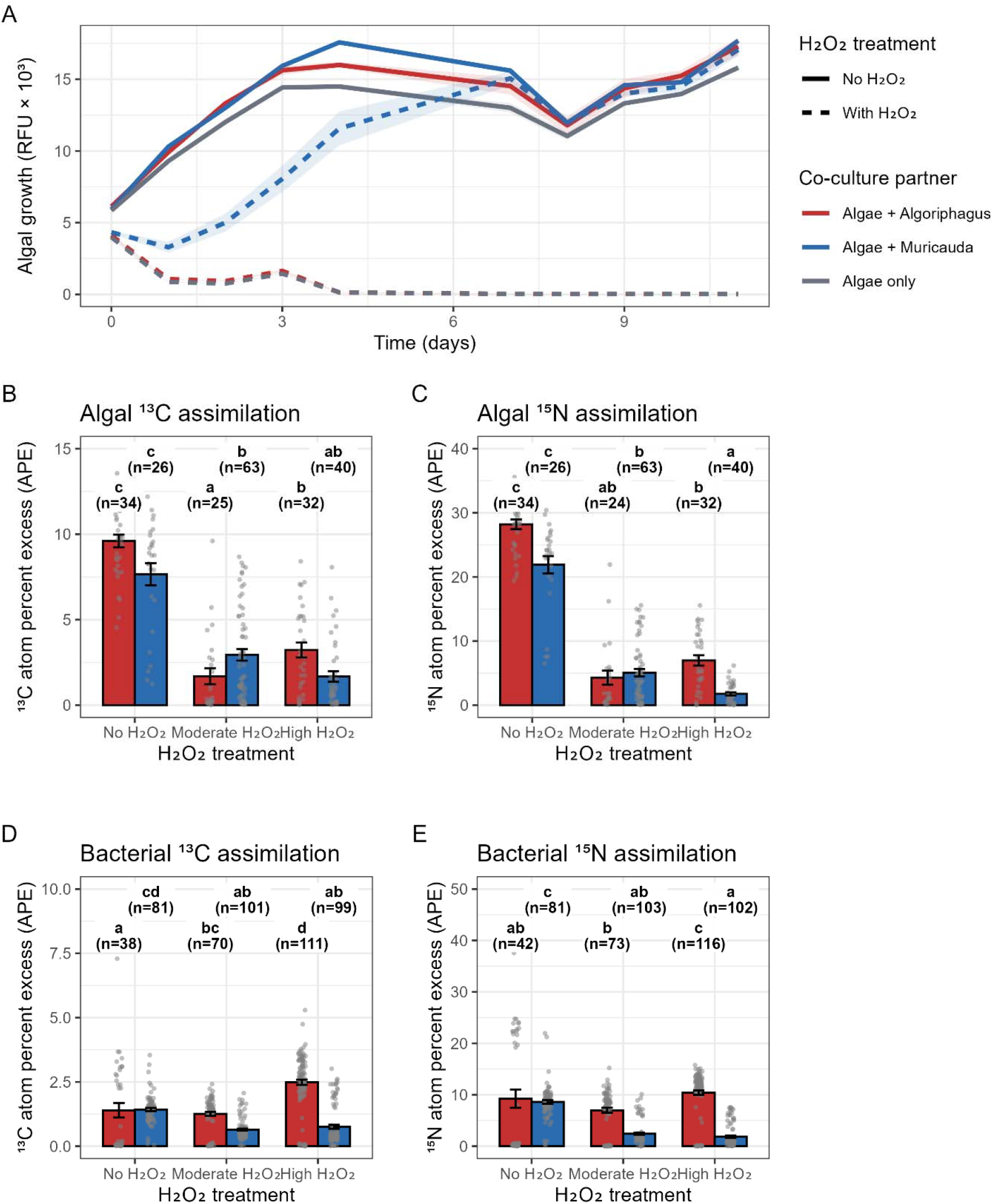
Effects of oxidative stress on algal and bacterial growth and metabolic activity. (A) Growth of *Phaeodactylum tricornutum* in monoculture and coculture with *Muricauda* or *Algoriphagus* under no H_2_O_2_ (solid lines) and H_2_O_2_ exposure (dashed lines). Lines show mean relative fluorescence (RFU × 10^3^); shaded ribbons show ± SEM. (B-C) Algal incorporation of inorganic ^13^C (B) and ^15^N (C) under no, moderate, and high H_2_O_2_ stress when grown alone or with bacterial partners. (D-E) Bacterial incorporation of algal-derived ^13^C (D) and dietary and algal-derived ^15^N (E) under the same treatments. Bars denote means ± SEM of single-cell nanoSIMS measurements. Different letters indicate statistically distinct groups (Tukey-adjusted comparisons following Type III ANOVA on log-transformed data; α = 0.05). Grey points represent individual NanoSIMS measurements from single cells. The value n represents the number of individual cells analyzed by NanoSIMS for each treatment group.

We then quantified algal assimilation of inorganic carbon and nitrogen under oxidative stress using nanoscale secondary ion mass spectrometry (nanoSIMS). *P. tricornutum* was grown alone or in coculture with *Muricauda* or *Algoriphagus* and exposed to no, moderate, or high H_2_O_2_. The concentrations of added H_2_O_2_ were lower than what is required to kill axenic *P. tricornutum* cultures because we were interested in investigating algal-bacterial interactions of ROS helpers and non-helpers without the algal host being killed. However, these concentrations (hundreds of micromolar) are higher than what is detected in ocean waters (1, 7, 9). High H_2_O_2_ involved multiple additions over time to maintain higher ROS concentrations. As expected, H_2_O_2_ exposure significantly suppressed algal assimilation of both inorganic carbon (^13^C-bicarbonate) and nitrogen (^15^N-nitrate) relative to no-addition controls (Figure 1B-C, Type III ANOVA, *P* < 0.001), consistent with impaired metabolic and physiological activity under oxidative stress (10,11). Algal assimilation depended on bacterial partner identity in an H_2_O_2_ concentration-dependent manner, with significant microbe × H_2_O_2_ interactions for both carbon and nitrogen, whereas microbial partner identity alone had no detectable effect. Under moderate oxidative stress, diatom cells cocultured with *Muricauda* showed significantly higher ^13^C assimilation than those cocultured with *Algoriphagus* (Tukey HSD, P < 0.01), while ^15^N assimilation did not differ between cocultures. This selective increase in carbon assimilation is consistent with a carbon-based reward model, in which helper-mediated H_2_O_2_ removal supports continued host carbon fixation and the production of labile carbon that can be transferred to the helper bacterium.

Under high oxidative stress, this pattern shifted and ^13^C assimilation did not differ significantly between cocultures, while ^15^N assimilation was significantly higher in diatoms cocultured with *Algoriphagus* than with *Muricauda* (Tukey HSD, P < 0.01), suggesting that high oxidative stress differentially affected carbon and nitrogen metabolism in diatom cells.

Next, we quantified the transfer of *P. tricornutum*-derived nutrients to *Muricauda* and *Algoriphagus*. Bacterial incorporation of algal-derived carbon (^13^C) and nitrogen (^15^N) varied with both bacterial identity and oxidative stress (Figure 1D-E), with significant effects of microbe, H_2_O_2_ treatment, and their interaction for both isotopes (Type III ANOVA, *P* < 0.01). Under both moderate and high oxidative stress, non-helper *Algoriphagus* consistently incorporated more algal-derived carbon and nitrogen than helper *Muricauda* (Tukey-adjusted *P* < 0.01), indicating greater access to phytoplankton-derived nutrients under ROS exposure. These results show that helper bacteria gained fewer resources from diatom hosts under oxidative stress, whereas non-helper bacteria benefited from more resource acquisition, consistent with opportunistic exploitation of host stress and damage.

We further investigated whether bacteria obtained *P. tricornutum*-derived nutrients primarily from exudates (compounds released by intact cells) or from lysates (compounds released upon cell damage). We incubated *Muricauda* and *Algoriphagus* monocultures in *P. tricornutum*-derived exudates and lysates and monitored growth by flow cytometry. The data showed that *Muricauda* grew with both exudate and lysate as the organic matter source, with higher cell yields with exudate. *Algoriphagus*, in contrast, exhibited little growth with exudate but substantial growth with lysate (Figure 2A). These data suggest that while both *Algoriphagus* and *Muricauda* could access nutrients from damaged or lysed *P. tricornutum* cells, only *Muricauda* was able to utilize extracellular organic resources released from healthy intact hosts.

**Figure 2.**
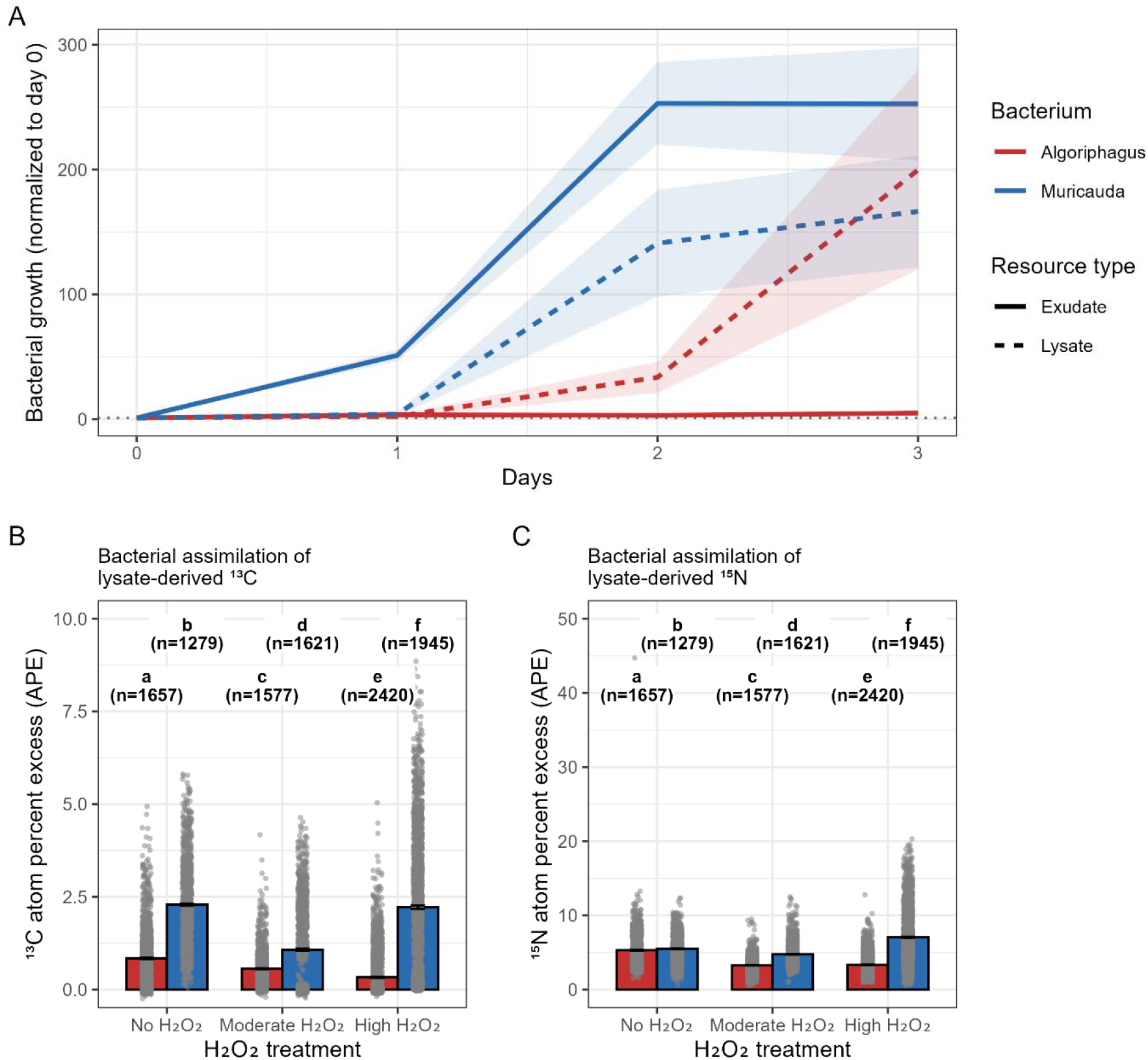
Oxidative stress differentially alters bacterial access to phytoplankton-derived resources. (A) Growth of *Muricauda* and *Algoriphagus* in *P. tricornutum* exudates or lysates (normalized to day 0). Lines indicate means; shaded regions show ± SEM. (B-C) Bacterial assimilation of lysate-derived ^13^C (B) and ^15^N (C) under no, moderate, and high H_2_O_2_. Bars show means ± SEM. Different letters denote significantly different groups based on Dunn’s post-hoc tests with Benjamini-Hochberg FDR correction following Kruskal-Wallis tests (α = 0.05). Grey points represent individual NanoSIMS measurements from single cells. The value n represents the number of individual cells analyzed by NanoSIMS for each treatment group.

We then evaluated how oxidative stress influenced bacterial assimilation of lysate-derived nutrients with a second stable isotope labeling experiment. Isotope labeled *P. tricornutum* lysate was generated and monocultures of helper *Muricauda* and non-helper *Algoriphagus* were incubated with this labeled substrate as the sole organic nutrient source under no, moderate, and high H_2_O_2_ concentrations. Both ^13^C and ^15^N assimilation varied significantly across all treatments (Kruskal-Wallis: ^13^C H(5) = 3469.99, *P* < 0.0001; ^15^N H(5) = 3158.27, *P* < 0.0001; Figure 2B-C), with multiple significant pairwise differences identified by Dunn’s post-hoc tests (FDR-adjusted *P* < 0.05). Under oxidative stress, *Muricauda* displayed partial metabolic resilience, with lysate carbon incorporation declining only under moderate ROS stress. Nitrogen incorporation was also reduced under moderate stress and was highest under high stress. On the other hand, *Algoriphagus* exhibited metabolic suppression with nutrient incorporation decreasing by up to ∼98% (^13^C) and 30% (^15^N) under oxidative stress. These contrasting responses revealed distinct physiological responses to oxidative stress between helper and non-helper bacteria.

The patterns of nutrient transfer, differential growth on exudates and lysates, and stress-dependent assimilation observed here suggest that oxidative stress reshaped phytoplankton-bacteria interactions by shifting the dominant mechanism of nutrient release from exudation by intact, metabolically active algal cells to leakage and lysis from damaged cells. Helper bacteria such as *Muricauda* appear primarily adapted to capitalize on exudate-derived substrates produced by healthy phytoplankton, whereas non-helper bacteria such as *Algoriphagus* preferentially exploit lysate-derived nutrients generated during host stress or mortality. This functional partitioning of strategies links oxidative stress directly to changes in nutrient availability and suggests that oxidative stress may select for different bacterial lifestyles in the phycosphere by altering the form in which host-derived nutrients are released.

Together, these results indicate that ROS scavenging by *Muricauda* was highly context dependent. Although it conferred metabolic resilience in monoculture, it became a costly strategy in coculture, where protecting the algal host reduced *Muricauda*’s incorporation of host-derived carbon and nitrogen. The observed reduction in isotope incorporation reflects a short-term metabolic tradeoff, which may be offset over longer timescales by the persistence of a healthy host. Conversely, *Algoriphagus* primarily benefited when oxidative stress damaged the diatom cells by gaining increased access to lysate-derived substrates. The exudate-lysate growth assays further suggest that these costs and benefits were tied to resource availability. *Muricauda* appeared to benefit primarily when *P. tricornutum* remained healthy, gaining access to exudate-derived substrates that are only produced under low oxidative stress. By reducing extracellular H_2_O_2_, *Muricauda* promoted algal survival and thereby maintained this exudate supply. *Algoriphagus*, conversely, benefited when *P. tricornutum* cells were damaged, gaining access to lysate-derived nutrient pools. The contrasting use of exudates and lysates provides a functional basis for context-dependent cooperation and suggests a process favoring partner fidelity, whereby *Muricauda*’s benefits are linked to maintaining a healthy host, whereas *Algoriphagus* thrives on algal stress or mortality.

Although the specific compounds mediating this helper-beneficiary interaction remain unidentified, likely candidates include small metabolites such as vitamins, organic acids, and amino acids, as well as more complex molecules such as extracellular polymeric substances (EPS;12,13). Future work characterizing phytoplankton exudate composition under oxidative stress and examining interactions among multiple bacterial partners will be valuable for understanding how ROS shape microbial community assembly and function in the phycosphere.

Although our findings were demonstrated here in a simplified phytoplankton-bacterium system, they suggest a broader conceptual framework in which oxidative stress drives algal-bacterial interactions by regulating the balance between cooperative and exploitative nutrient acquisition strategies. We propose that under low to moderate oxidative stress, helper bacteria can maintain host health and access exudate-derived resources, promoting partner fidelity between phytoplankton and their bacterial associates. However, under frequent or intense oxidative stress, the capacity of helpers to protect their hosts may be exceeded, resulting in increased host damage and the release of lysate-derived resources that can be exploited by opportunistic taxa. Testing this framework across diverse phytoplankton-bacterium systems and environmental contexts will be essential for determining the extent to which oxidative stress shapes microbial interactions and nutrient cycling in aquatic ecosystems.

## Author contributions

N.A. and X.M. conceived and developed the study. N.A., C. S., N. M., K. R., C. R., and X. M. gathered the data and conducted the analyses. K. R. and C. R. contributed to sample preparation. N.A. led the writing of the manuscript. R. K. S. and P. K. W. contributed critically to the analyses and writing. All authors edited the manuscript and approved the final version.

## Data availability

Data that support the findings of this study are available in the supplementary information. Source data are provided with this paper.

## Supplementary Information

### Supplementary Methods

#### Bacterial coculture experiments with Phaeodactylum tricornutum

To assess the effects of bacterial coculture on *Phaeodactylum tricornutum* growth under oxidative stress, coculture experiments were conducted using 22 individual bacterial isolates. The bacterial strains used were: *Alcanivorax* sp. EA2, *Algoriphagus* sp. ARW1R1, *Arenibacter* sp. 7G5Y1, *Devosia* sp. EAB7W2, *Henriciella* sp. 6ES, *Hoeflea* sp. G46H50, *Limnobacter* sp. G6BE5s, *Marinobacter* sp. Pt3-2, *Mesorhizobium* sp. DxT, *Muricauda* sp. ARW1Y1, *Muricauda* sp. 7G5W, *Oceanicaulis* sp. 13A, *Pseudonocardia* sp. DxW, *Pusilimonas* sp. EA3, *Rhodophyticola* sp. 6CLA, *Roseibium* sp. 13C1, *Roseobacter* sp. N2s, *Stappia* sp. ARW1T, *Sulfitobacter* sp. N5S, *Tepidicaulis* sp. EA10, *Thalassospira* sp. 13M1, and *Yoonia* sp. 4BL. Axenic *P. tricornutum* cultures and *P. tricornutum* cocultured with individual bacterial strains were grown under control conditions (no H_2_O_2_ addition) or in the presence of H_2_O_2_. Cultures were grown in 250 µL of enriched artificial seawater (ESAW) in 96-well plates. For H_2_O_2_ treatments, H_2_O_2_ was added to final concentrations of 195 µM or 326 µM, while control cultures received no H_2_O_2_ addition. Algal growth was monitored daily for 10–11 days using chlorophyll autofluorescence as a proxy for bulk biomass (excitation/emission: 440/680 nm), measured with a BioTek Cytation 5 multimode plate reader. Biological triplicates of all treatment and control samples were incubated with shaking (90 rpm, 22°C;12 h:12 h, light:dark; 3500 lux illumination).

#### Coculture isotope experiment

To investigate the impact of coculture with bacteria on metabolic activity under H_2_O_2_ stress, cocultures of *Phaeodactylum tricornutum* with *Algoriphagus* sp. ARW1R1 and *Muricauda* sp. ARW1Y1 were grown under H_2_O_2_ stress conditions. Cultures (5 mL) were incubated in individual wells of a 12-well plate. All cultures were grown in enriched artificial seawater (ESAW) medium supplemented with 2 mM ^13^C-sodium bicarbonate and 0.828 mM ^15^N-nitrate (Cambridge Isotope Laboratories, Tewksbury, MA). Two H_2_O_2_ treatment conditions were applied, a double-spike treatment, in which H_2_O_2_ was added to a final concentration of 100 µM at 0 and 24 h post-inoculation, and a four-spike treatment, in which H_2_O_2_ was added to a final concentration of 100 µM at 0, 24, 32, and 45 h post-inoculation. These H_2_O_2_ additions were lower than the concentrations that lead to axenic *P. tricornutum* cell death, specifically to examine stressful but not killed cultures incubated with a ROS helper and a non-helper. A control treatment with no H_2_O_2_ addition was also included. Biological triplicates of all treatment and control samples were incubated at 22 °C with agitation at 90 rpm under 3500 lux illumination on a diurnal 12 h:12 h light:dark cycle for 52 h. Algal growth was monitored throughout the incubation period using a BioTek Cytation 5 Multimode Reader. After incubation, samples for NanoSIMS analysis were fixed by adding 500 µL of 37% formaldehyde to 5 mL of culture, vortexed briefly, and stored at 4 °C until filtration. Fixed samples were centrifuged at 3,000 rpm for 5 min prior to filtration onto 0.2 µm, 25 mm type GTTP white polycarbonate filters (Millipore, Burlington, MA). Cells retained on the polycarbonate filters were washed three times with sterile, filtered Milli-Q water to remove residual labeled media. The washed filters were stored at 4 °C in the dark until NanoSIMS analysis.

#### Generation of isotopically labelled Phaeodactylum tricornutum lysate

Isotopically labelled lysates were generated by sonicating ^13^C and ^15^N-labelled *P. tricornutum* cells. Labelled *P. tricornutum* cultures were produced by incubating axenic *P. tricornutum* in ESAW supplemented with 10 mM ^13^C-sodium bicarbonate and 2 mM ^15^N-nitrate (as described above) for six days, and transferred again into isotope labeled media to maximize labeling. Isotope-labelled *P. tricornutum* cells were lysed by sonication using the microtip on a VWR Misonix Sonicator 3000 at 3 for 30 second bursts, repeated three times. The sonicator dial was subsequently set at 4 for 60 second bursts, repeated five times. Samples were kept on ice for two minutes between each sonication throughout the sonication process to prevent overheating. The resulting lysate was then passed through a 0.22 µm filter to generate cell debris free lysate. Lysates were stored at -80 °C until use.

#### Labeled lysate experiment

To investigate the impact of exogenous *Phaeodactylum tricornutum*-derived nutrient supplementation on heterotrophic bacterial H_2_O_2_ degradation, heterotrophic bacteria were grown in isotopically labelled lysates under conditions of H_2_O_2_ stress. Two H_2_O_2_ treatment regimes were applied, a double-spike treatment, in which 100 µM H_2_O_2_ was added at 0 and 3 h post-inoculation, and a four-spike treatment, in which 100 µM H_2_O_2_ was added at 0, 3, 19, and 27 h post-inoculation. A control sample with no H_2_O_2_ addition was also included. Biological triplicates of all treatment and control samples were incubated at 24 °C for 43 h. Bacterial growth was monitored by flow cytometry. Samples (100 µL) were fixed with 2 µL of 25% glutaraldehyde, vortexed, and stored at -80 °C until analysis using an Attune NxT flow cytometer (Thermo Fisher Scientific Inc.). After incubation, samples for NanoSIMS analysis were collected by adding 200 µL of 37% formaldehyde to 2 mL of culture, vortexing, and storing at 4 °C until filtration. Fixed samples were filtered as described above and analyzed by NanoSIMS.

#### NanoSIMS analytical conditions

Samples collected on filters were prepared for NanoSIMS analysis by cutting each filter into 1/8-size wedges using sterile scissors or scalpels. Filter sections were mounted onto aluminum sample disks using conductive carbon tabs (Ted Pella #16084-6) and coated with approximately 5 nm of gold to enhance surface conductivity. Isotopic imaging was performed using a CAMECA NanoSIMS 50 instrument at Lawrence Livermore National Laboratory. Analyses were conducted using a ^133^Cs^1^ primary ion beam operated at 1.5 pA, corresponding to an approximate beam diameter of 150 nm at an impact energy of 16 keV. Prior to data acquisition, each analysis area was pre-sputtered with a higher primary beam current (90 pA) to a depth of ∼60 nm to remove surface contamination and achieve sputtering equilibrium. For isotope imaging, the primary beam was rastered over 20 × 20 μm areas with a dwell time of 1 ms per pixel. Each field of view was scanned repeatedly for 19-30 cycles, generating images with a resolution of 256 × 256 pixels. The secondary ion mass spectrometer was tuned to a mass resolving power of approximately 7000 (1.5× corrected), sufficient to resolve relevant isobaric interferences. During acquisition, secondary electron images and quantitative secondary ion images were collected simultaneously for ^12^C□ □, ^13^C^12^C□, ^12^C^14^N□, and ^12^C^15^N□ on individual electron multipliers operated in pulse-counting mode. Isotopic ratios were calculated as ^13^C^12^C□/^12^C□ □ (equal to 2 × ^13^C/^12^C) and ^12^C^15^N□/^12^C^14^N□ (equal to ^15^N/^14^N). All nanoSIMS datasets were initially processed using the software package L’Image (Larry Nittler, Arizona State University) to correct for detector dead time and image drift. Corrected ion images were then used to generate ^13^C/^12^C and ^15^N/^14^N ratio images, which reflect the incorporation of ^13^C- and ^15^N-labeled substrates into cellular biomass. Individual cells were identified based on ^12^C^14^N□ images, and regions of interest (ROIs) corresponding to single cells were manually delineated. For each ROI, isotopic ratios were calculated on a cycle-by-cycle basis, and mean values and standard errors were determined. To quantify substrate incorporation, we calculated the excess fraction of cellular carbon and nitrogen derived from the labeled substrates (C_net_ and N_net_, respectively). These values were computed using the isotope fraction measured in each target cell (*F*_*c*_), the isotope fraction measured in killed controls representing initial cellular enrichment (*F*_*kc*_), and the isotope fraction of the added substrate (*F*_*s*_). ROI-level data were exported for further statistical analysis in R, where mean values and standard errors of C_net_ and N_net_ were calculated for each treatment.

#### Statistical Analysis

All statistical analyses were performed in R (v4.4.2). NanoSIMS isotope incorporation datasets (^13^C and ^15^N atom percent excess, APE) for coculture experiments were analyzed using linear models with microbial partner, H_2_O_2_ treatment, and their interaction as fixed factors. Because APE distributions were right-skewed and included very low values, data were log-transformed after addition of a small positive offset. Effects were assessed using Type III ANOVA, and when significant, differences among groups were evaluated using Tukey-adjusted pairwise contrasts (emmeans v1.11.2.8). Compact letter annotations indicate significance groupings (α = 0.05). For lysate-assimilation datasets a non-parametric approach was applied since distributions deviated from normality and variances were heterogeneous. Differences among microbe × H_2_O_2_ groups were first tested using Kruskal-Wallis rank-sum tests, followed by Dunn’s post-hoc tests with Benjamini-Hochberg false discovery rate (FDR) correction for pairwise comparisons. Full statistical outputs are provided in the Supplementary Information.

## Figure legends

Figure 1. Effects of oxidative stress on algal and bacterial growth and metabolic activity. (A) Growth of *Phaeodactylum tricornutum* in monoculture and coculture with *Muricauda* or *Algoriphagus* under no H_2_O_2_ (solid lines) and H_2_O_2_ exposure (dashed lines). Lines show mean relative fluorescence; shaded ribbons show ± SEM. (B-C) Algal incorporation of inorganic ^13^C (B) and ^15^N (C) under no, moderate, and high H_2_O_2_ stress when grown alone or with bacterial partners. (D-E) Bacterial incorporation of algal-derived ^13^C (D) and dietary and algal-derived ^15^N (E) under the same treatments. Bars denote means ± SEM of single-cell nanoSIMS measurements. Different letters indicate statistically distinct groups (Tukey-adjusted comparisons following Type III ANOVA on log-transformed data; α = 0.05).

Figure 2. Oxidative stress differentially alters bacterial access to phytoplankton-derived resources. (A) Growth of *Muricauda* and *Algoriphagus* in *P. tricornutum* exudates or lysates (normalized to day 0). Lines indicate means; shaded regions show ± SEM. (B-C) Bacterial assimilation of lysate-derived ^13^C (B) and ^15^N (C) under no, moderate, and high H_2_O_2_. Bars show means ± SEM. Different letters denote significantly different groups based on Dunn’s post-hoc tests with Benjamini-Hochberg FDR correction following Kruskal-Wallis tests (α = 0.05).

## Supplementary Figures

**Figure S1.**
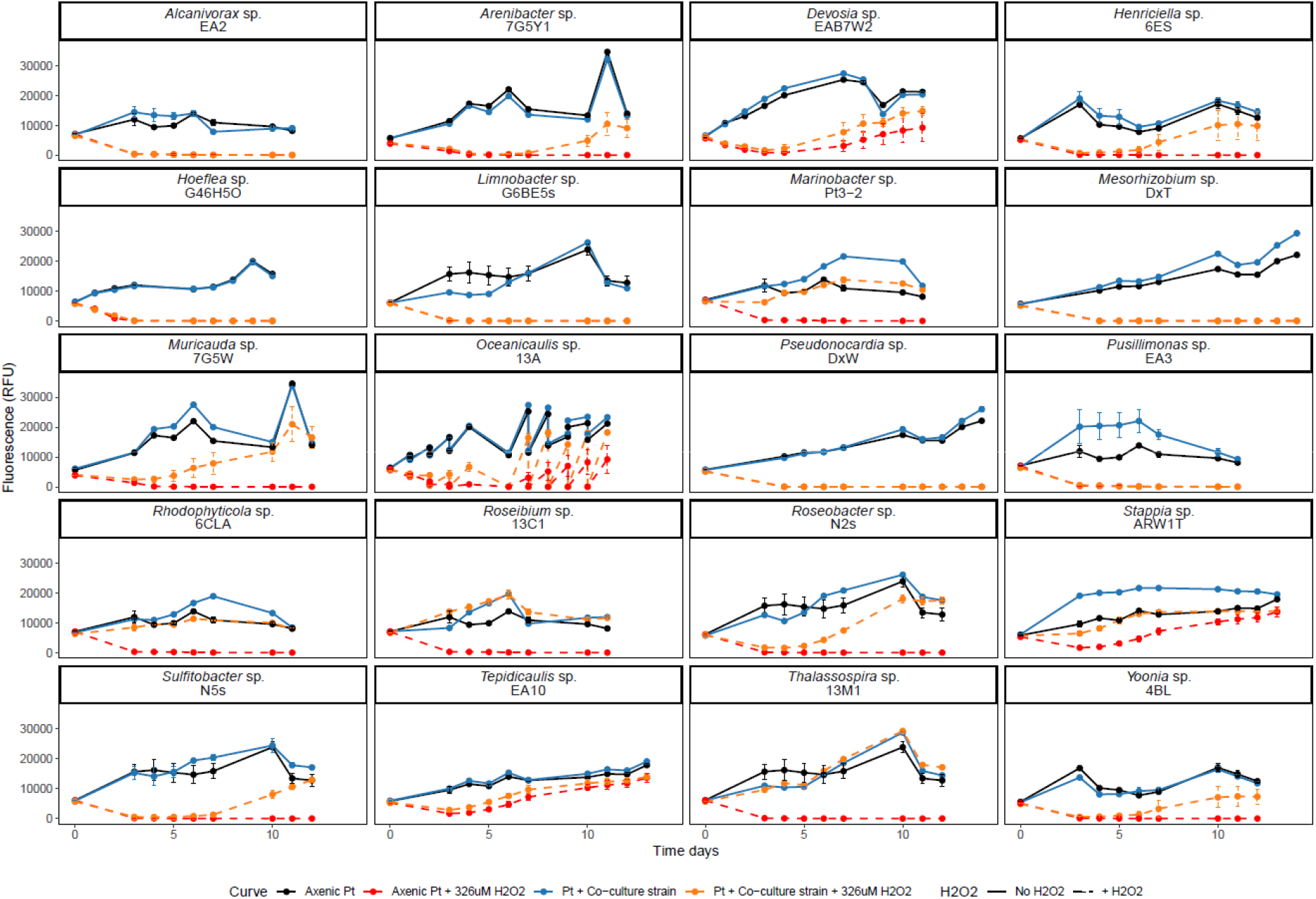
Growth dynamics of axenic *Phaeodactylum tricornutum* and coculture strains under oxidative stress. Growth curves show mean fluorescence (RFU) over time for axenic *Phaeodactylum tricornutum* (Pt) and *P. tricornutum* cocultured with individual bacterial strains under control conditions (no H_2_O_2_) or in the presence of H_2_O_2_. Axenic Pt is shown in black (control) and red (+H_2_O_2_), while coculture conditions are shown in blue (control) and orange (+H_2_O_2_). Solid lines indicate control conditions, and dashed lines indicate H_2_O_2_ treatment. Points represent mean values calculated from n = 3-4 biological replicates, and error bars denote the standard error of the mean (SEM). Each panel represents a single coculture strain. Strain names are shown in the panel titles.

**Figure S2.**
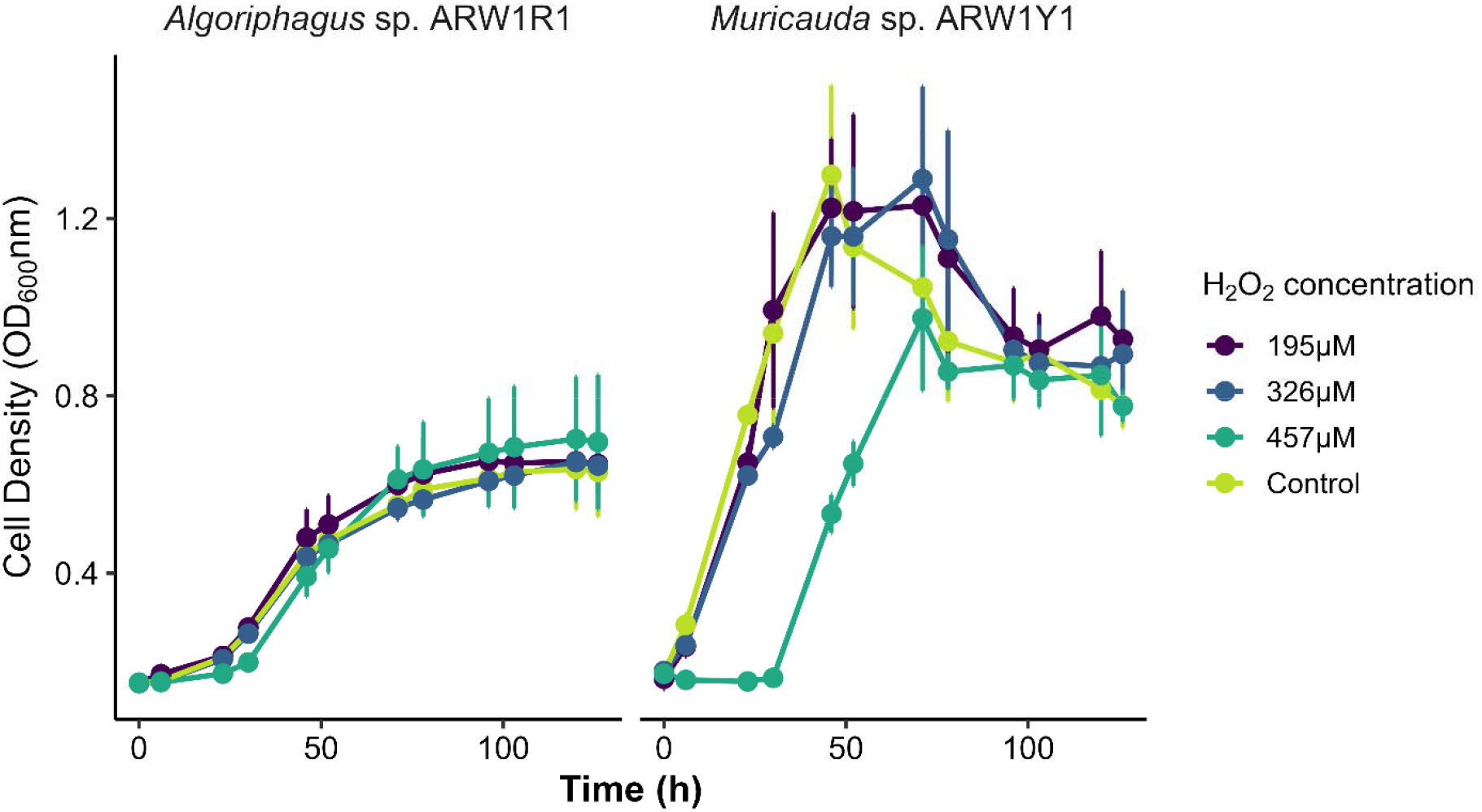
Effect of hydrogen peroxide on the growth of *Algoriphagus* and *Muricauda*. Cell density of *Algoriphagus* (left panel) and *Muricauda* (right panel) was monitored over time following exposure to increasing concentrations of hydrogen peroxide. Error bars represent standard deviation (SD) of three biological replicates.

**Figure S3.**
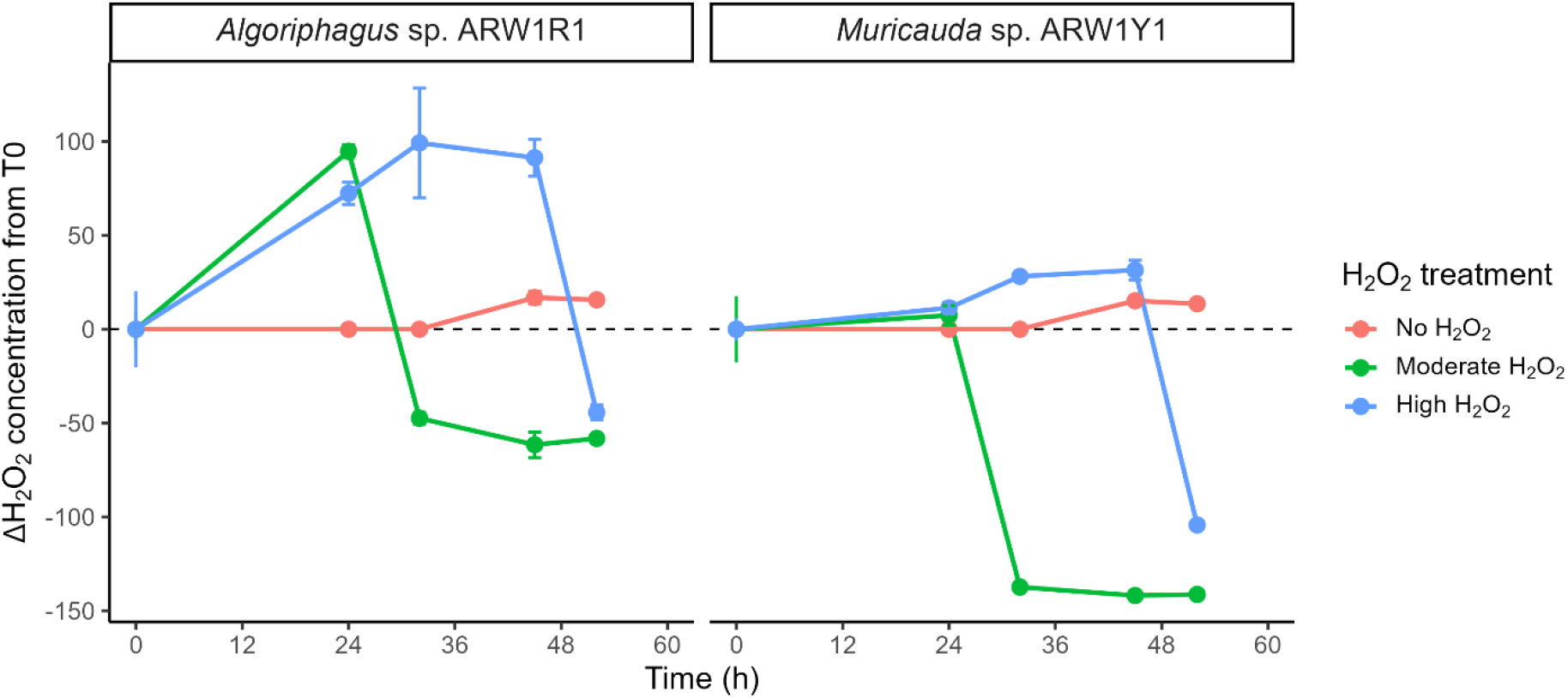
Temporal changes in H_2_O_2_ concentration in *Phaeodactylum tricornutum* cocultures with *Algoriphagus* and *Muricauda*. H_2_O_2_ concentrations were measured over time in cocultures containing either *Algoriphagus* (left panel) or *Muricauda* (righ panel) exposed to no H_2_O_2_, moderate H_2_O_2_, or high H_2_O_2_ treatments. Values are shown as the mean ± SD of replicate measurements. To account for differences in initial H_2_O_2_ concentrations among treatments, data were normalized by subtracting the mean concentration at time zero (T0) for each treatment from all subsequent measurements. All treatments begin at a value of zero, positive values indicate H_2_O_2_ accumulation and negative values indicate depletion relative to the initial condition.

## References

1. Lesser MP. (2006). Oxidative stress in marine environments: biochemistry and physiological ecology. Annu Rev Physiol 68: 253–278.

2. Diaz JM, Plummer S. (2018). Production of extracellular reactive oxygen species by phytoplankton: past and future directions. J Plankton Res 40: 655–666.

3. Morris JJ, Rose AL, Lu Z. (2022). Reactive oxygen species in the world ocean and their impacts on marine ecosystems. Redox Biol 52: 102285.

4. Cirri E, Pohnert G. (2019). Algae–bacteria interactions that balance the planktonic microbiome. New Phytol 223: 100–106.

5. Fischer WW, Hemp J, Valentine JS. (2016). How did life survive Earth’s great oxygenation? Curr Opin Chem Biol 31: 166–178.

6. Hünken M, Harder J, Kirst GO. (2008). Epiphytic bacteria on the Antarctic ice diatom Amphiprora kufferathii Manguin cleave hydrogen peroxide produced during algal photosynthesis. Plant Biol 10: 519–526.

7. Morris JJ, Johnson ZI, Szul MJ, Keller M, Zinser ER. (2011). Dependence of the cyanobacterium Prochlorococcus on hydrogen peroxide-scavenging microbes for growth at the ocean’s surface. PLoS One 6: e16805.

8. Morris JJ, Lenski RE, Zinser ER. (2012). The Black Queen Hypothesis: evolution of dependencies through adaptive gene loss. mBio 3: e00036–12.

9. Zinser ER. (2018). The microbial contribution to reactive oxygen species dynamics in marine ecosystems. Environ Microbiol Rep 10: 412–427.

10. Robles-Rengel R, Florencio FJ, Muro-Pastor MI. (2019). Redox interference in nitrogen status via oxidative stress is mediated by 2-oxoglutarate in cyanobacteria. New Phytol 224: 216–228.

11. Weenink EF, Matthijs HC, Schuurmans JM, Piel T, van Herk MJ, Sigon CA, et al. (2021). Interspecific protection against oxidative stress: green algae protect harmful cyanobacteria against hydrogen peroxide. Environ Microbiol 23: 2404–2419.

12. Brisson V, Swink C, Kimbrel J, Mayali X, Samo T, Kosina SM, Thelen M, Northen TR, Stuart RK. (2023). Dynamic Phaeodactylum tricornutum exometabolites shape surrounding bacterial communities. New Phytol 239: 1420–1433

13. Orellana LH, Francis TB, Ferraro M, Hehemann JH, Fuchs BM, Amann RI. (2022). Verrucomicrobiota are specialist consumers of sulfated methyl pentoses during diatom blooms. ISMEJ 16:630–41.

